# Deep Learning Detection and Classification of Red Blood Cells: Towards a Universal Dataset

**DOI:** 10.1101/2025.09.18.677048

**Authors:** A. Amrani, I. Caridi, B. Kaoui

## Abstract

We evaluate emerging machine learning models for pattern recognition, focusing on the YOLOv11 architecture for detecting and classifying red blood cell shapes. Our analysis targets two characteristic morphologies observed under flow: *slipper* and *parachute*. A key challenge in this task is the development of a robust and diverse dataset. To address this, we employ synthetic image generation using a cut-and-paste approach, introducing variations in cell overlap and arrangements of microfluidic channels disposition to alleviate data scarcity and reduce cross-dataset bias. We generate these datasets with U-Net and Cellpose segmentation models, and rigorously assess YOLOv11 performance on two benchmarks: (i) a controlled dataset for evaluating classification accuracy, and (ii) a challenging, visually heterogeneous dataset for assessing generalization. Results show that the model achieves high precision for distinct cell types in controlled settings, but exhibits reduced performance on the unseen dataset, highlighting a trade-off between specialized accuracy and broad applicability in complex microscopy scenarios.

## I. INTRODUCTION

Red Blood Cells (RBCs) are the most abundant cell type moving through the vessels of the cardiovascular system. They play the crucial role of delivering oxygen to organs and tissues throughout the body, while also removing carbon dioxide. Their equilibrium shape depends strongly on their mechanical properties and on the environment they are in. Certain adopted shapes can indicate pathological conditions [13]. Previous experimental studies have classified RBC shapes primarily using static images [17]. Recent advances in microfluidics and high-speed imaging now make it possible to capture emerging RBC shapes under dynamic flow conditions. For example, RBCs in flow can be broadly categorized into two main shapes: slipper (asymmetric) and parachute (symmetric). The slipper shape is elongated and asymmetric, while the parachute shape - also known as the croissant shape - has a folded configuration with a concave rear [12]. These deformations help RBCs pass through microscopic blood capillaries, where efficient gas exchange occurs between the RBC membrane and surrounding tissues. The shape of RBCs influences both the local flow patterns and the macroscopic rheology of blood. It can also serve as a diagnostic indicator for disorders such as malaria. Distinctive RBC forms, such as slipper and parachute, are observed under flow both *in vivo* (in the microvasculature) and *in vitro* (in microfluidic chips). Efficient, real-time classification of these flow-dependent shapes, and others, can provide valuable insights into RBC mechanical properties and related pathologies. Our goal is to create a universal labeled data set of RBC images, focusing mainly on slipper and parachute shapes, and to enable automated shape detection using the YOLO framework. Detection and classification of RBCs under flow conditions remains an active area of interest in medical imaging research, particularly with the growing adoption of Artificial Intelligence (AI) methods. To the best of our knowledge, only two key studies have addressed this problem in a systematic manner.

Kihm *et al*. [15] were the first to apply convolutional neural networks (CNNs) for the automatic classification of experimentally captured RBC shape images, tackling the inherent subjectivity and time demands of manual classification. In their approach, the image pre-processing pipeline began by standardizing all images to a 90*×* 90 pixel format, ensuring consistency across the dataset. A Tukey window with *α* = 0.25 was then applied to create a smooth fade-out towards the microchannel walls, thereby reducing the effect of irregular light refraction from channel edges, which is a common source of noise that can degrade CNN training and performance. To further enhance robustness, the image intensities were mapped to the full 8-bit range by saturating the bottom and top 1% of the pixel values. This adjustment increased the effective signal dynamic range and normalized intensity profiles, improving stability against variations in illumination. Unlike conventional binary classification approaches, their AI workflow was based on a regression-style CNN architecture that produced floating-point outputs. This design choice reflected the observation that RBCs in flow often exhibit a large number of “indefinite shapes” or transitional morphologies, which cannot be reliably assigned to fixed categorical classes. The network was trained to detect two main stable shapes, slippers and croissants, while also including an auxiliary “sheared croissant” class to boost classification precision, despite its ambiguous morphology in two-dimensional imaging. The supervised training process used a dataset of 4, 000 manually labeled RBC images, which was doubled to 8, 000 through mirroring. CNN used Root Mean Square Error (RMSE) as the loss function and used stochastic gradient descent with momentum for optimization. Training was intentionally limited to 10 epochs to avoid overfitting, and validation loss was monitored as the primary performance metric. Importantly, the dataset encompassed images recorded under varying flow strengths, lighting conditions, and focal planes (including slightly defocused images), ensuring that the trained model was robust to optical misalignments and experimental variability. The continuous CNN output was then post-processed using Gaussian function fitting to define statistically derived thresholds. This enabled the automated construction of unbiased RBC shape state diagrams, which showed strong agreement with those obtained through manual classification. Liang *et al*. [18] proposed a method for assessing RBC deformability that markedly improved measurement sensitivity through refined shape classification. Their AI-based workflow advanced classification granularity by employing YOLOv5 to simultaneously detect and categorize multiple RBCs within a single image. The system classified RBCs into six distinct morphological types: Parachute (symmetrical, centered), semi-Parachute (slightly asymmetrical), semi-Slipper (transitional state with rear opening), Slipper (bulging front with narrow tail), semi-Rolling (rigid, shrunken oval), and Rolling (undeformed disk). Model training was based on thousands of manually annotated images, resulting in a final dataset of 2, 415 labeled cells across the six classes. YOLOv5s was chosen for its computational efficiency, completing training in approximately 1.5 hours over 216 epochs, and achieving an accuracy of 0.84. The trained model operated with a confidence threshold of 0.6 and an Intersection over Union (IoU) threshold of 0.5, enabling reliable detection and classification of multiple cells per image.

We would like to emphasize an important, non-negligible point: in both of the aforementioned studies, the authors analyzed images that they had themselves acquired. The generalization of their proposed AI models to new, unseen, and unknown datasets has not yet been tested. This is precisely the gap we address in our present work. To overcome data scarcity, we employed an automatic data construction process, described in Section II D and detailed in Section III B 3. Our contributions to this research field are as follows: i - Construction of a Red Blood Cell dataset containing thousands of cells acquired, under different conditions, by various research groups, with the aim of creating a universal dataset, ii - Evaluation of the generalization capabilities of the YOLO framework, trained on this dataset, by testing it on a new unseen dataset with substantially different characteristics, iii - Discussion of detection and classification performance, with particular attention to factors that influence these results. The code and data sets are available on GitHub [1].

## II. MODELS AND NUMERICAL METHODS

### A. Literature survey

Classification of the shapes of RBCs at rest, at equilibrium, has been extensively studied and reported in the literature. Various classification criteria have been proposed considering abnormalities in shape (e.g. echinocytes, ovalocytes, spherocytes), size (microcytes, macrocytes), color (hypochromic, hyperchromic), membrane composition and mechanical properties, and surface markers (e.g., blood typing). In contrast, the categorization of RBC shapes in images obtained under flow conditions, an out-of-equilibrium situation, has been less thoroughly investigated. Under such conditions, two main morphologies are typically identified: slippers and parachutes, corresponding to shapes adopted by RBCs in flow regimes similar to those in blood vessels. The parachute shape is generally observed at low flow velocities and particularly at higher reduced volumes (*τ*), as illustrated in Figure 1. As flow velocity increases, RBCs tend to transition from this symmetric parachute shape to an asymmetric slipper-shaped morphology (see Figure 1). This slipper shape predominates at higher flow velocities and lower reduced volumes, suggesting that increased flow rates and reduced cell inflation favor the formation of slipper morphologies, as discussed in Ref. [12].

**FIG. 1:**
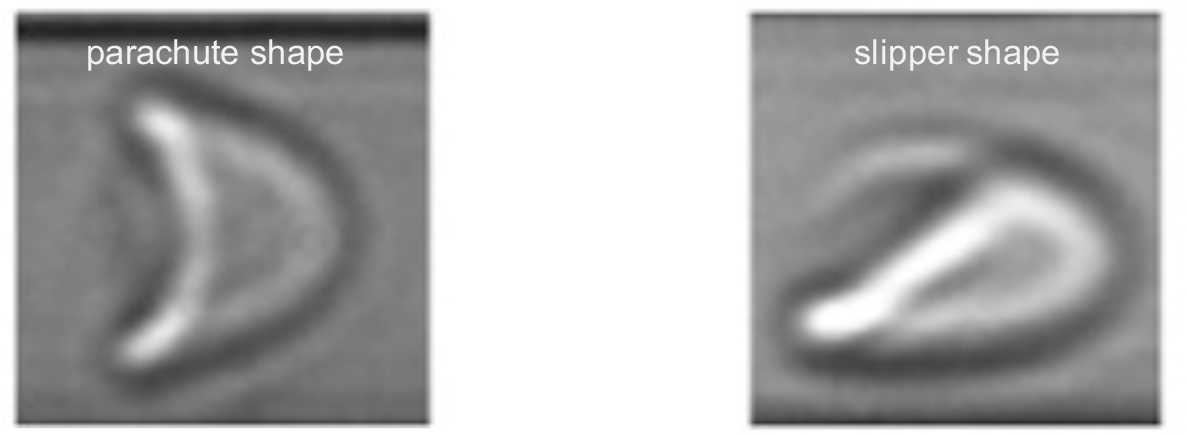
Experimentally observed steady shapes of Red Blood Cells (RBCs) under flow: symmetric parachute shape (left panel) and asymmetric slipper shape (right panel), adapted from Ref. [15].

We conducted a survey of potential dataset providers to find out about the available RBC image datasets that we could use in our study.

- **Zenodo** [38]: Zenodo, founded and run by researchers at CERN in collaboration with the European Commission’s OpenAIRE project, is an open-access data repository for sharing research data. It serves as a general-purpose platform, inviting researchers from all disciplines worldwide to submit and deposit their work openly, including datasets, software, and other digital artifacts. In accordance with open science standards, Zenodo does not impose restrictions on data format. For individual uploads, researchers can include up to 50, GB of data, distributed across a maximum of 100 files. For larger projects, Zenodo allows researchers and EU-funded initiatives to request a one-time quota increase for a single record, raising the limit to 200, GB. To optimize the uploading and downloading process—particularly for datasets with multiple files, compressing them into a single archive is strongly recommended.
- **Kaggle** [16]: Kaggle is an online community for data scientists and machine learning engineers. It was founded in 2010 and is run by Google. The platform’s diverse services include hosting data science competitions, providing access to public datasets, and offering a cloud-based environment for data science and artificial intelligence education. Kaggle allows users to upload individual datasets of up to 200, GB. For file structuring, up to 50 top-level files are supported per dataset. Additionally, there is a cumulative storage limit of 200, GB for all private datasets associated with a single user account.
- **BioStudies** [3]: BioStudies is an open database launched in 2018. It is dedicated to the systematic deposition of data from biological studies. Located within the European Bioinformatics Institute (EMBL-EBI), it serves as a centralized repository for a wide range of biological data types, with the aim of enhancing data accessibility and re-usability for the scientific community. Due to its focus on inclusive biological study data, submission guidelines emphasize accurate data presentation and the provision of rich metadata, ensuring that submitted datasets retain their value and contextual background. Researchers with very large datasets are encouraged to contact the BioStudies team at the time of submission.
- **GitHub** [10]: GitHub is a popular cloud-based platform best known for version control and collaborative software development. It provides a space for users to store, share, and collaborate on code projects. Although not designed for hosting large datasets, it can be used to share small datasets or image file collections related to code. For single files uploaded via the web interface, GitHub imposes a limit of 100, MB. To handle larger files, GitHub offers Git Large File Storage (LFS), which supports the management of files exceeding this limit. While it is technically possible to store up to 5, GB in a repository, standard GitHub repositories are generally recommended to remain under 1, GB in total size.
- **Figshare** [8]: Figshare is an open-access repository where researchers can save and share their research outputs, including datasets, figures, videos and other materials. Figshare provides all users with a limited amount of free storage upon registration, while institutional accounts often have higher storage allowances. The specific dataset limits depend on the type of account—free or institutional. It generally allows individual file uploads of several gigabytes, with overall storage limits determined by the account subscription.
- **National Institutes of Health (NIH) Data Commons** [21]: NIH Data Commons is an initiative of the National Institutes of Health that provides a cloud environment for hosting, sharing, and analyzing biomedical data, including omics and imaging datasets. It aims to accelerate biomedical discoveries by improving accessibility of information to the research community. Accessing and sharing data within the NIH Data Commons generally requires adherence to specific protocols, data standards, and collaborative agreements established for the NIH-supported research community. Dataset restrictions and sharing policies are typically defined within the framework of specific NIH programs and data collections hosted in the Commons.
- **University and Hospital Databases:** Several universities and hospitals maintain internal image databases to support their research activities. These resources are generally not accessible to external researchers unless accessed through direct contact with the institution.

There are other repositories focused on biomedical data, such as PhysioNet (primarily for physiological signals), The Cancer Imaging Archive (for cancer imaging), and OpenNeuro (for neuroimaging), which are not relevant to our project.

To identify the relevant data sets of interest to our study in the selected providers, we used a targeted automatic search strategy that used a combination of specific keywords. This approach aimed to maximize the retrieval of datasets containing images of RBCs that exhibit slipper and parachute morphologies under flow conditions. The following terms were combined in various search queries across each platform:

- **Cell type focus:** “Red Blood Cell”, “RBC”, or “Erythrocyte” are used to ensure that the search focuses on the correct biological cell type,
- **Image modality:** Terms such as “shape”, “morphology”, “image”, or “microscopy” have been included to prioritize datasets containing visual data,
- **Target shapes:** The keywords “slipper” and “parachute” are used together to specifically target datasets that exhibit these morphologies. “croissant” can also be used instead of “parachute”, as many articles use the two terms synonymously,
- **Flow conditions:** Terms such as “flow”, “fluid dynamics”, or “microfluidics” are incorporated to prioritize datasets that show the behavior of RBCs under flow conditions,
- **Dataset Request:** The term “dataset” is included in providers like GitHub, where there may be other file types on the platform.

Some of other possible combinations investigated are: “red blood cell slipper parachute”, “RBC image slipper parachute”, “RBC flow microscopy”, “erythrocyte shape microfluidics”, “RBC image classification flow”, “RBC slipper parachute dataset”, “RBC shape in flow images”, “red blood cell morphology flow”.

### B. Dataset impact on learning

A significant challenge that can impede the performance and reliability of AI models is the presence of cross-dataset bias. This phenomenon arises when AI models are trained or evaluated on datasets that possess differing statistical properties, a common occurrence given the diverse ways in which visual data is collected and annotated. Such disparities between datasets can lead to a trained model exhibiting high accuracy on its training data, while suffering a significant decline in performance when applied to new and unseen data from a different source. A particularly salient aspect of cross-dataset bias in vision detection models stems from either the variations in camera characteristics across different datasets used for training or the content differences in datasets. The latter is a widely known problem, notably since the publication of the *Name That Dataset!* experiment [35]. The goal was to proove that a linear Support Vector Machine can differentiate the dataset from which input images belongs to. The results highlight that each dataset, even when targeting the same goal (ie. car images for car detection in the cited paper), has a *signature* based on the content of the dataset. This implies that a model trained on a dataset will show worse performance when tested on a dataset that has another signature. The notion of signature evoke the particularity of a dataset: [35] takes the example of car detection datasets, each having a particular manner of taking picture of cars (some datasets contain mainly pictures of cars taken from the side, while others may have picture taken by the front).

Another major obstacle in dataset creation is camera characteristics. When a difference occurs in camera settings, as pixel size and focal length, between the training dataset and testing dataset, the model might show worse performance. The impact of such parameters has been studied in [19], where the same neural network topology was trained on different datasets, each with their own pixel size. The result show that the generalization of the model differs based on the pixel size: the same model, trained on the same content but with different pixel size, will perform differently on the same unseen dataset. The authors emphasize that each machine learning problem might need a different AI model architecture and dataset needs. It is thus not possible to call for particular camera settings. Other parameters as exposure have an impact on model’s learning, but these can be mitigated with pre-processing steps.

Further investigations shows that a trained model may recognize a object differently based on the camera used to photograph the object [4]. Camera lenses have different field of view, making the object appear larger or smaller. This first characteristic can slightly change the object category given by the trained model during evaluation (ie. from “cups” to “coffee mug” in the cited study). Taking a step further, evaluating the trained model on the same object, using different cameras with the same field of view also shows differences in the predicted category on the same objects.

The above features highlight the impact of the learning material used on the performance of the AI vision model. From this, two choices can be made to train an AI model. On the one hand, if the evaluation conditions are known (ie. if the user knows precisely the camera used in production, the distance to objects, the backgrounds), the training dataset should be composed of related images. However, if the evaluation conditions are unknown, it is preferable to design a robust model by maximizing the diversity of images used in training, both on the object side (the same object pictured under different points of view with a diverse set of backgrounds) and on the material side (using different camera characteristics).

### C. Image processing techniques

Image processing techniques are applied to create a final dataset that includes all relevant identified images. To augment the available data and facilitate different analytical approaches, we also create an intermediate dataset. It consists of images with well-centered RBCs, organized into two distinct folders corresponding to their classified shapes: slipper and parachute. The final data set contains images with multiple RBCs. This dataset, whose construction method is described in Section II D, is structured into two folders: an “image” folder containing the final images, and a “label” folder. Within the “label” folder, each image has a corresponding text file detailing the labels and bounding box coordinates of all RBCs present in that image, following the format (label, *x* center, *y* center, height, width), where position and size values are within the [0, 1] interval, normalized with respect to image height and width. The image processing tasks are primarily implemented in Python, utilizing several libraries, including PyTorch (version 2.7.0) for the U-Net segmentation model, OpenCV (version 4.11.0.86) for image manipulation tasks such as cropping and resizing, and scikit-learn (version 1.6.1) for the DBSCAN clustering algorithm.

#### 1. U-Net model

U-Net is a convolutional neural network architecture specifically designed for biomedical image segmentation. Its architecture is characterized by a contracting path (encoder) that progressively downsamples the input image, capturing contextual information through a series of convolutional layers, pooling operations, and feature map reductions. This contracting path is followed by a symmetric expanding path (decoder) that gradually upsamples the lower-resolution feature maps, combining them with high-resolution features from the contracting path via skip connections. These skip connections enable the network to propagate fine-grained details from the early layers to the later layers, allowing for precise localization of the segmented regions. The final layers of the expanding path produce a segmentation map with pixel-wise class predictions [29]. Due to its ability to learn effectively from limited annotated data through strong data augmentation and its robust localization capabilities, U-Net has become a reference architecture for a variety of biomedical image segmentation tasks. Indeed, the U-Net architecture, along with its variations and pre-trained versions, has demonstrated strong generalization across a wide range of biomedical imaging modalities, achieving high performance in segmenting objects in microscopy images (e.g., cells, nuclei, organelles), radiology images, and ophthalmic imaging. The U-Net used in this work is implemented in PyTorch [20].

#### 2. CellPose

Cellpose is a deep learning-based method specifically developed for cellular segmentation from microscopy images [30]. Its primary objective is to serve as a generalist algorithm capable of precisely segmenting cells from a wide range of image types without requiring model retraining or parameter adjustments for each new image type. This addresses a significant challenge in cell segmentation, where methods typically require new human-labeled images to achieve good performance on different data types. The core innovation of Cellpose is its use of an auxiliary representation that transforms cell masks into two image maps of the same size as the original image. These maps represent horizontal and vertical spatial gradients that point toward the center of each cell. Unlike classical methods such as the watershed algorithm, which rely directly on image intensity values and can struggle with cells that have inhomogeneous marker distributions, Cellpose’s vector fields are designed so that all pixels within a cell converge to a fixed point representing the cell center. This vector flow representation forms a single smooth topological basin, making segmentation more robust to variations in cell intensity. A neural network is trained to predict these horizontal and vertical gradients, as well as a binary map indicating whether a pixel is inside or outside a region of interest. The core of the Cellpose architecture is based on the U-Net convolutional network, with several modifications to improve performance: replacing standard building blocks with residual blocks, doubling the network depth, and using additive skip connections instead of concatenation to reduce the number of parameters. The training dataset included fluorescently labeled cytoplasm and membranes, DAPI-stained nuclei in separate channels, brightfield microscopy images, and even non-microscopy images of repeated objects such as fruits and rocks, hypothesized to aid generalization. This diverse training set allows the generalist Cellpose model to generalize far more widely than methods trained on specialized datasets from a single laboratory or image type. We used the Cellpose segmentation model to extract the bounding polygons of the cells, which were then cut from their original images and pasted onto a generated background, as described in III B 2 and III B 3.

#### 3. Density-Based Spatial Clustering of Applications with Noise

DBSCAN (Density-Based Spatial Clustering of Applications with Noise) is a density-based clustering algorithm that groups densely populated points (high-density regions) and identifies points in low-density areas as noise. Unlike partitioning algorithms such as k-means, DBSCAN does not require a predefined number of clusters. The algorithm is based on two parameters: epsilon (*ϵ*), the radius around a data point within which to search for neighbors, and *minPts*, the minimum number of data points within this radius required for a point to be considered a core point. Core points lie inside a cluster, while border points are within the neighborhood of a core point, but have fewer than *minPts* neighbors themselves. Points that are neither core points nor border points are classified as noise. By gradually expanding clusters from core points to density-connected points, DBSCAN can effectively identify clusters of arbitrary shape, is robust to noise, and is therefore a valuable tool for data exploration, anomaly detection, and pre-processing steps such as outlier removal in image processing pipelines [7]. The DBSCAN implementation used in this work is provided by scikit-learn [24].

#### 4. Crop Technique

The image cropping technique involves selecting and extracting a specific rectangular region of interest from a larger image, discarding the portions outside this boundary. Cropping is commonly used to focus analysis on a particular object or area within an image, remove irrelevant background information, or create a dataset of isolated objects from images containing multiple instances.

#### 5. Resizing technique

The image resizing technique involves altering the dimensions of an image, either increasing (upsampling) or decreasing (downsampling) its width and height. This process is often used to standardize the input size for machine learning models, reduce the computational cost of processing large images by decreasing their resolution, or prepare images for display or storage at specific dimensions. Various interpolation methods, such as nearest neighbor, bilinear, or bicubic interpolation, can be used during resizing to determine the pixel values in the new image grid based on the original image data, each offering different trade-offs between computational complexity and the smoothness or sharpness of the resulting image. The resizing technique used in this work is implemented in PyTorch [36].

### D. Dataset creation with cut-and-paste

To address the scarcity of images containing multiple objects, new training images can be synthetically generated by first extracting foreground objects (the cells) from known images, and then pasting them onto different backgrounds. This creates a dataset of images, each containing multiple cells. This technique, known as the cut-and-paste method [6], has been evaluated and shown competitive results compared to models trained solely on real images (*−*10% to *−*0.9% in average precision at IOU 0.5 for training on cut-and-paste images versus real images for everyday objects in indoor environments). It should be noted that models trained on both synthetic and real images outperformed models trained only on real images for the selected task in the reference study. Such a dataset could be used in future work. However, directly employing this technique may exacerbate the domain gap. As in any AI framework, the model can suffer from distribution mismatches between training and testing images (i.e., training images exhibiting characteristics different from testing images) [9]. In addition, the cut-and-paste method may multiply domain gaps, since both the foreground and background source domains (cells pasted on backgrounds forming the training images) may differ from the foreground and background target domains (testing images), thereby enlarging the gap between the total training and testing domains (real images from microfluidic devices). For example, if the background source-to-target domain gap is smaller than the foreground source-to-target domain gap (meaning the training background images are relatively similar to the target, but the foreground objects are not), the AI model might be well-trained on the background but struggle with the foreground objects. This can lead to many variations of foreground objects being incorrectly classified as background, resulting in a high false rejection rate [37]. A solution is to reduce the gap between the source domain of objects (training cells) and the target domain of objects (cells in the testing data), while also minimizing the background source-to-target domain gap [37]. The cited study overcame this issue by diversifying the foreground objects through image transformations (altering color characteristics), while shrinking the background training domain using Gaussian blur and grayscale transformation. This approach showed improved performance compared to the default cut-and-paste method presented in [6]. Domain inspection is thus an important step in the cut-and-paste method. Nonetheless, we can assume that our use of this method does not introduce a massive domain gap, as the source domain of the background is already limited (noisy grayscale backgrounds, with noise and grayscaling being solutions found in [37]). Only the foreground domain gap of the cells may be significant if the cells used in the testing data differ substantially. Moreover, analysis of the training images using *H*-divergence, as in [2], should be conducted in future work to validate this assumption. Further information on the implementation of the cut-and-paste method is presented in section III B 3.

### E. YOLO model

The second goal of our study is the assessment of the performance of the YOLO (You Only Look Once) model, using both zero-shot and fine-tuning approaches, on the dataset we created. YOLO is a real-time object detection system that revolutionized the field by framing object detection as a single regression problem, outputting class probabilities and bounding boxes directly from an image in a single evaluation. Its architecture typically involves a single convolutional neural network that divides the input image into a grid. For each cell of the grid, the network simultaneously predicts several bounding boxes and a per-box confidence score, quantifying the likelihood of object presence and the accuracy of the box prediction. In addition, for each predicted box, the network predicts a probability distribution over the object classes. By processing the entire image at once, YOLO achieves much faster inference rates than methods based on region proposal steps and is well-suited for real-time scenarios [27]. Later variants of YOLO have emerged, introducing architectural improvements and training practices to increase accuracy and address multi-object detection challenges in various domains [11]. The version used in this work is YOLOv11. YOLOv11 is the latest release in Ultralytics’ series of real-time object detectors. Building on the success of previous YOLO models, YOLOv11 introduces substantial architectural and training improvements, making it a competitive architecture for a wide variety of computer vision tasks. Key enhancements include a stronger feature extraction backbone and neck, leading to more precise object detection and better performance on demanding tasks. Optimized for speed and efficiency, YOLOv11 offers faster processing times without compromising the trade-off between accuracy and performance. Additionally, YOLOv11 is designed for seamless deployment across various environments, from edge devices to cloud and NVIDIA GPU-enabled platforms, supporting applications such as object detection, instance segmentation, image classification, pose estimation, and oriented object detection [14].

### F. Computational Environment

The system was developed and tested on both Windows and Linux platforms. Data preparation, including image segmentation and dataset construction, was carried out on an Intel Core i7-8565U CPU (base frequency 1.80 GHz). Model training and evaluation of the YOLO framework were performed on an NVIDIA GH200 Grace Hopper Superchip, which substantially improved training speed and inference efficiency. Although the trained YOLO model can operate on lightweight hardware, the use of a dedicated graphics processing unit is strongly recommended when working with large datasets in order to reduce inference time. The code and data sets are available on GitHub [1].

## III. RESULTS AND DISCUSSION

### A. Literature search results

The results are reported in Table I for all dataset providers. The initial investigation did not yield relevant datasets on most platforms. Specifically, although BioStudies and Figshare provide some video content, no datasets containing static images labeled with slipper or parachute shapes were identified.

**TABLE I:** Summary of Dataset Search Results.

| Dataset Provider | Result | Note |
| --- | --- | --- |
| Zenodo | No results | Filter selected for resources types: dataset; and for file type: png, jpg, zip. |
| Kaggle | No results | Research performed in the dataset section. |
| BioStudies | Videos in rectangular microchannel [31];<br>Synthetic videos in microchannel [26];<br>Reconstruction of slipper and parachute shapes [32] [22] |  |
| GitHub | No results | Research on web as not presented a specific dataset section |
| Figshare | Synthetic videos in microchannel [5] |  |
| National Institutes of Health (NIH) Data Commons | No results |  |
| University Databases | For the following university there are some datasets with slipper/parachute shapes: Universität des Saarlandes (Germany) [15] [23]; Università degli studi di Napoli Federico II (Italy) [34] [33]; Singapore University of Technology and Design (Singapore) [18] | The authors were contacted directly and they shared their data |

We have successfully identified and accessed several valuable datasets needed for our study thanks to academic partners:

- **Singapore University of Technology and Design (SUTD) -** The dataset [18] is accessible for download via the ResearchGate platform [28]. This dataset contains 5000 grayscale images (50 *×* 100 pixels) categorized into six classes: parachute (732 images), slipper (391), rolling (390), semi-parachute (311), semi-slipper (201), and semi-rolling (234),
- **Universität des Saarlandes (UdS) -** The first dataset [15] contains 90 *×* 90 grayscale images of well-centered RBCs, classified into three categories: slipper, sheared, and parachute. The dataset is organized into two folders: an input data folder containing 1336 slipper, 1110 sheared, and 644 parachute images, and a training data folder containing 3000 slipper, 2000 sheared, and 3000 parachute images. The second dataset [23] contains 92 *×* 92 grayscale images of well-centered RBCs, divided into three classes: slipper (81 images), other (58), and croissant (76). Regarding this dataset, we also obtain the corresponding original videos divided in blood donors, three in total, and two pressure values used to extract the RBC shapes (100 mbar and 1000 mbar). The videos are not classified.
- **Università degli studi di Napoli Federico II (UniNa) -** The provided dataset contains nine folders classified on the velocity of the cell, where the minimum is 0.12 cm*/*s and the maximum is 1.56 cm*/*s; each folder contains a variable number of images. The dataset is not classified [33].

### B. Dataset creation

#### 1. Extraction of cells for datasets with one RBC

Using the dataset from the SUTD [18], we focus on four RBC morphologies: parachute, slipper, semi-parachute, and semi-slipper. To construct our final dataset with three categories, we introduce a combined class, sheared, which merges the semi-parachute and semi-slipper images. Since the RBCs are not consistently centered, a pre-processing pipeline is required. We first segment each image to isolate individual cells, as their precise coordinates are not provided. An initial attempt with zero-shot object segmentation using YOLO failed to delineate cell boundaries reliably. Therefore, we employ a U-Net convolutional neural network, which is well-suited for biomedical image segmentation. Following segmentation, we crop each image based on the masks to obtain well-centered RBCs. Figure 2 illustrates an example of a segmentation mask and the corresponding cropped image.

**FIG. 2:**
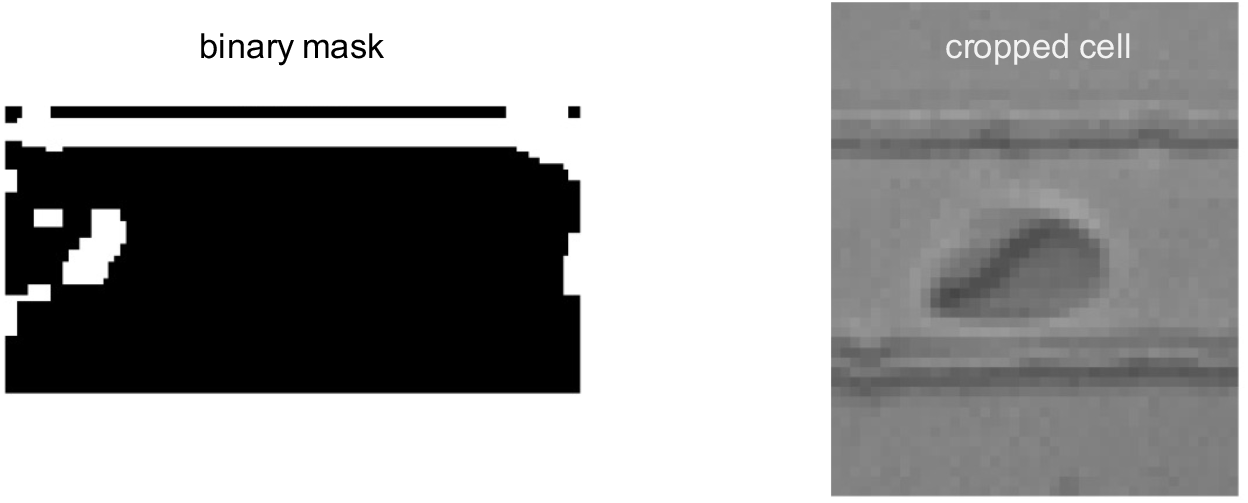
Example of a processed image based on data related to the work Ref. [18]: U-Net mask (left panel) and cropped image of a red blood cell with a slipper shape (right panel)

Since the cropped images vary in size, we resize all segmented cell images to a standardized dimension of 50 *×* 50 pixels. This standardization ensures consistent input dimensions for subsequent analysis. After resizing, we apply the DBSCAN algorithm to identify and remove potential outlier objects in the segmented images. This density-based clustering approach enables us to discard cases where the cell is not well-centered, as illustrated in Figure 3. The resulting dataset consists of slipper (248 images), sheared (464 images), and parachute (682 images). For the other datasets, the images had already undergone preprocessing. In particular, for the first dataset from UdS [15], we merged the training and test folders into a single dataset. After processing each dataset to contain well-centered RBC images labeled as slipper and parachute, we resized all images to 90 *×* 90 pixels to standardize dimensions across datasets. The resized images from all sources were then combined to create a unified dataset structured into three categories: slipper, sheared, and parachute. We applied the same preprocessing pipeline to the video datasets from UdS [23] and UniNa [33] universities. However, U-Net was unable to segment the cells in these cases. Consequently, these datasets were not included in the single-RBC datasets. The preprocessing of dataset [23] is detailed in the following section, while no preprocessing was required for the dataset of Ref. [33].

**FIG. 3:**
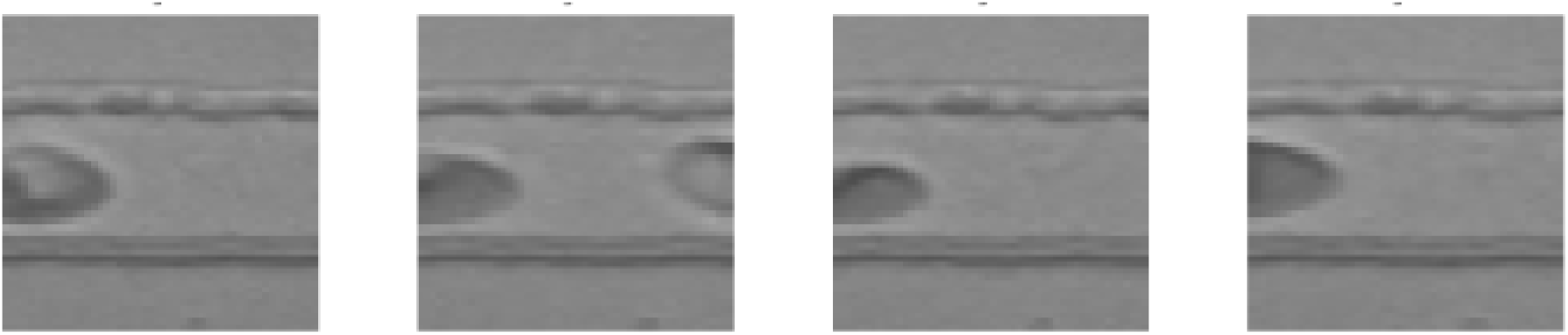
Examples of outliers, which are removed from the data using the DBSCAN algorithm. Example of outliers identified after the cell extraction starting from data related to the work in Ref. [18]

#### 2. Extraction of cells for datasets with multiple RBCs

This section considers datasets containing multiple cells per image. To apply the cut-and-paste method, an additional step is required to extract the cells without their surrounding background, as illustrated in Figure 4. We employ CellPose segmentation models on preprocessed images to obtain centered cell extractions. For the dataset [18], we use the cyto3 segmentation model with a diameter of 60 pixels. Since CellPose outputs may include outliers, we introduce filtering guardrails. First, because parachutes are typically centered in the channel, only centered segmentations are retained. This criterion does not apply to slippers, which are not consistently centered. For slippers, we apply a second guardrail, retaining only segmentations with a height-to-width aspect ratio greater than 0.5, corresponding to elongated shapes. This procedure yields 588 parachutes out of the 682 previously segmented, and 190 slippers out of the 248 previously segmented. For the dataset [15], we use the nuclei segmentation model with a fixed diameter of 60 pixels. For parachutes, the same centered guardrail is applied, while for slippers, only segmentations with an aspect ratio less than 3.5 are retained. This results in 2490 extracted parachutes out of 3000 and 2992 slippers out of 3000 in the dataset.

**FIG. 4:**
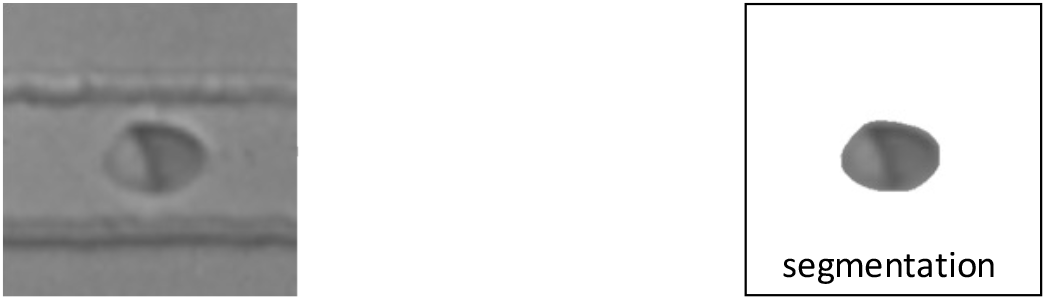
Segmentation of a RBC from an image of a well centered Red Blood Cell, data related to the work in Ref. [18]

For the videos [23], we use the nuclei segmentation model with a fixed diameter of 40 pixels. Images in this dataset contain many outliers, i.e., cells that are neither slippers nor parachutes. To address this, we implement a label inference algorithm to classify each cell extracted by CellPose. First, an upper bound on the number of pixels per extracted cell is set to 450 for slippers and 350 for parachutes, in order to exclude excessively large outliers. Since parachutes are characterized by a symmetric shape, we filter out cells where the absolute difference in pixel count between the top and bottom halves exceeds 5%. We further retain only cells with fewer pixels in the left half than in the right half, consistent with the left-to-right flow direction. For slippers, we apply complementary criteria. Cells are retained when the absolute difference in pixel count between the top and bottom halves is greater than 1%, reflecting the asymmetry caused by the tail. Additionally, only cells with more pixels in the left half than in the right half are kept, as the tail produces an empty region on the right side of the bounding box. Applying this process, we extract 204 parachutes and 1492 slippers. The comparatively low number of parachutes results from the strict pixel-based filtering (350 pixels), which is necessary to avoid inclusion of other cell shapes.

#### 3. Dataset construction

Once each cell is extracted from its source image, multiple cells can be pasted onto a background to form a final composite image. Since no dataset of microfluidic backgrounds is available, we generate synthetic backgrounds and artifacts, following the approach of [25], which uses synthetic cells and backgrounds to construct datasets for training neural networks. The parameters for generating backgrounds and artifacts are chosen as described in the supplementary materials of [25]. We summarize the process below.

Backgrounds are filled with a random gray shade (pixel values between 100 and 150), onto which Simplex and Gaussian noise are applied. Artifacts are then added: small black filled circles and thin hair-like black lines to mimic particles; larger straight white lines to reproduce the borders of the microfluidic device; and round cells deformed through elastic transformations to introduce distractors that are neither parachutes nor slippers. This process produces blurry gray images with artifacts resembling real microfluidic pictures, as illustrated in Figure 5.

**FIG. 5:**
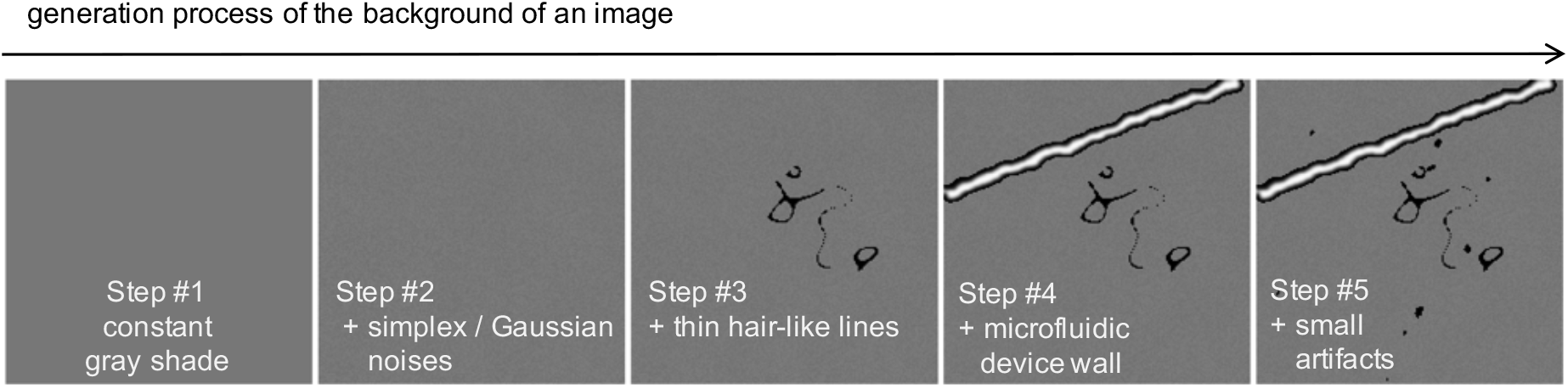
Generation process of the background of an image. From left to right: constant gray shade, Simplex and Gaussian noises, thin hair-like lines, microfluidic device wall, small artifacts.

After background generation, extracted parachutes and slippers, as well as synthetically generated cells from Ref. [25], are pasted onto the backgrounds. Each image contains at most four parachutes, four slippers, and four synthetic cells, each with random rotation, random placement on the background, and random scaling. Cells are positioned to avoid overlap with one another.

Finally, a noise function is applied to each image, chosen from *Salt-and-Pepper* noise, Gaussian noise, random Uniform noise, or no noise. Examples of the resulting noise effects are shown in Figure 6.

**FIG. 6:**
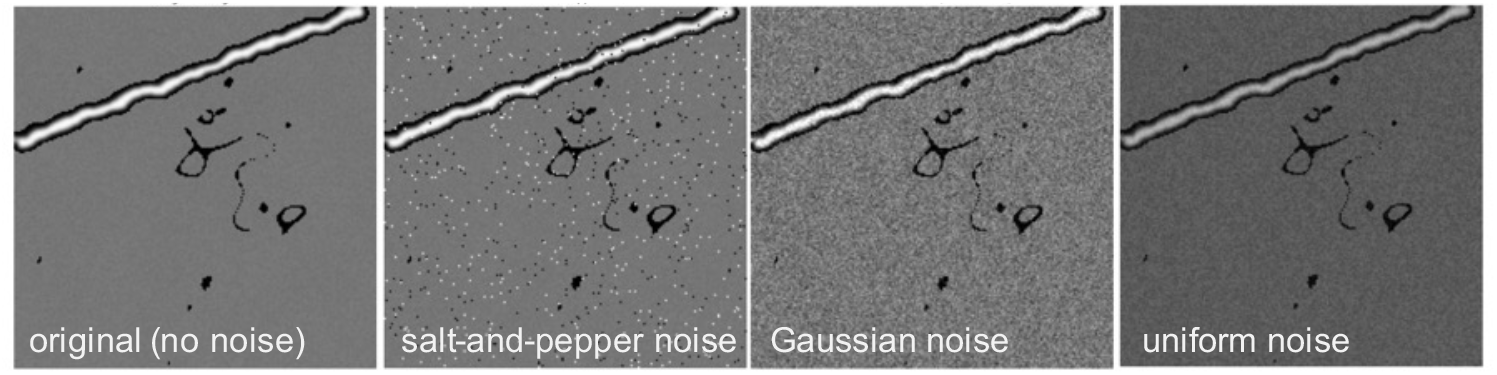
Example of noises. From left to right: original image (no added noise), Salt-and-Papper noise (sets random pixels to black and white), Gaussian Noise, Uniform Noise.

In order to prevent model performance from depending on generation parameters, we follow a factorial design across all relevant factors: the presence of a wall (yes/no), the presence of a thin line (yes/no), the presence of small black artifacts (yes/no), and the number of each type of cell (three or four). Taking the Cartesian product of these values yields an equally distributed set of parameters that is applied to each data split (training, validation, and testing). As a result, the final dataset contains balanced and independent distributions of cell counts and background elements. Examples of images from the generated dataset are shown in Figure 7.

**FIG. 7:**
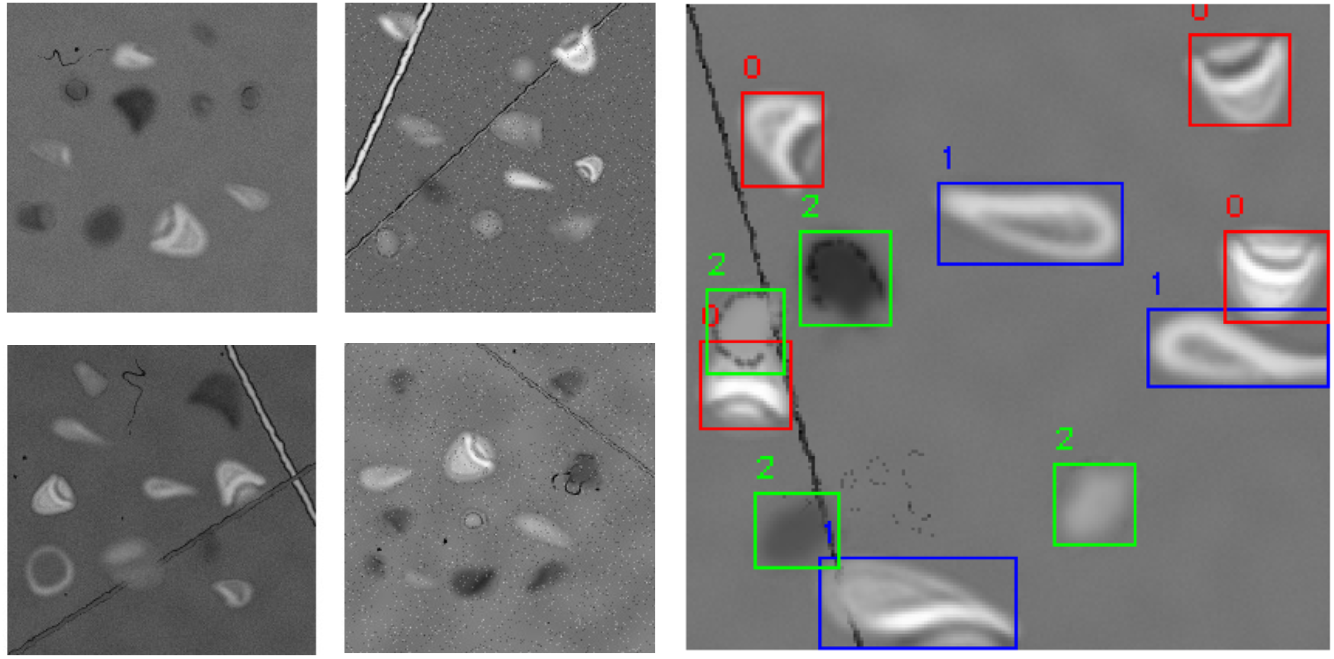
Example of final images. The dataset contains images with walls, images with artifacts and different types of noise. Each image has parachutes, slippers, and synthetically generated cells.

The dataset is created in two distinct ways with respect to cell arrangement: allowing possible superposition of cells and prohibiting superposed cells. In the former, cells may overlap when placed using the cut-and-paste method, whereas in the latter, overlaps are prevented. During dataset creation, it is important to ensure sufficient diversity to prevent the model from performing poorly in real-world scenarios. For example, if a dataset contains only images of cells moving from left to right, with all objects such as parachutes and slippers oriented identically, the model might struggle to identify these objects when presented with different orientations in production. To generalize the dataset and improve model performance, we varied the orientation and size of the cells. This approach helps the model recognize objects regardless of position or scale, leading to more reliable results in real-world applications. For each generated image, its characteristics (e.g., number of each cell type or presence of lines) were recorded and analyzed for linear correlation. The resulting correlation matrix is shown in Figure 8a. We conclude that no significant correlations are present in this constructed dataset.

**FIG. 8:**
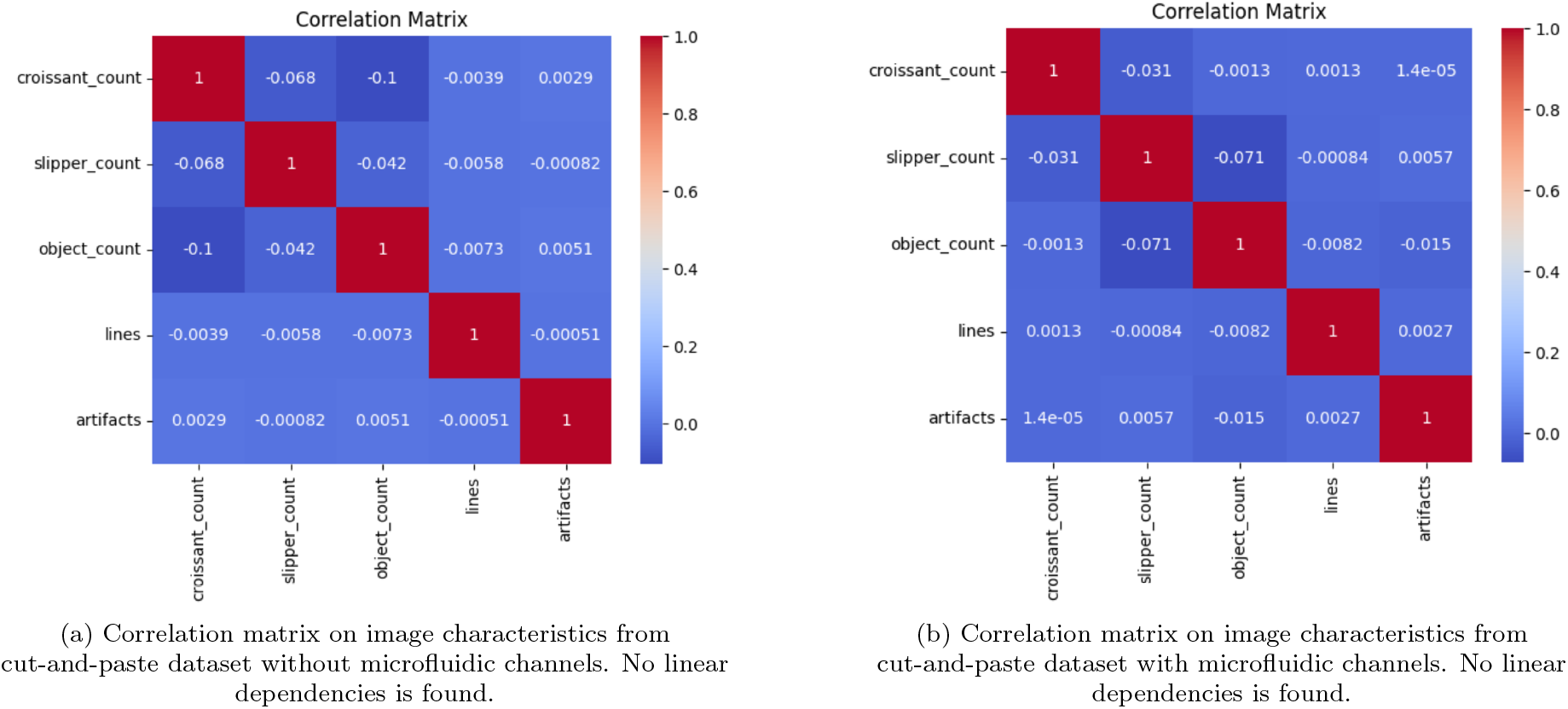
Confusion matrices for the generated cut-and-paste dataset without and with microfluidic channels.

Another dataset is created similarly to the previous one, but with the addition of a microfluidic channel to better mimic the acquisition of RBCs in flow dynamics. Images consist of a horizontal channel containing a variable number of cells. Cells are displayed with varying orientations to ensure robustness during training and are placed entirely within the channel. In this dataset, cells are not allowed to overlap, as overlaps do not occur in realistic microfluidic environments observed in our acquired datasets. Cells may touch, but they do not overlap. Examples of images from this generated dataset are shown in Figure 9. Correlations of image characteristics are presented in Figure 8b, and again, no significant linear correlations are observed.

**FIG. 9:**
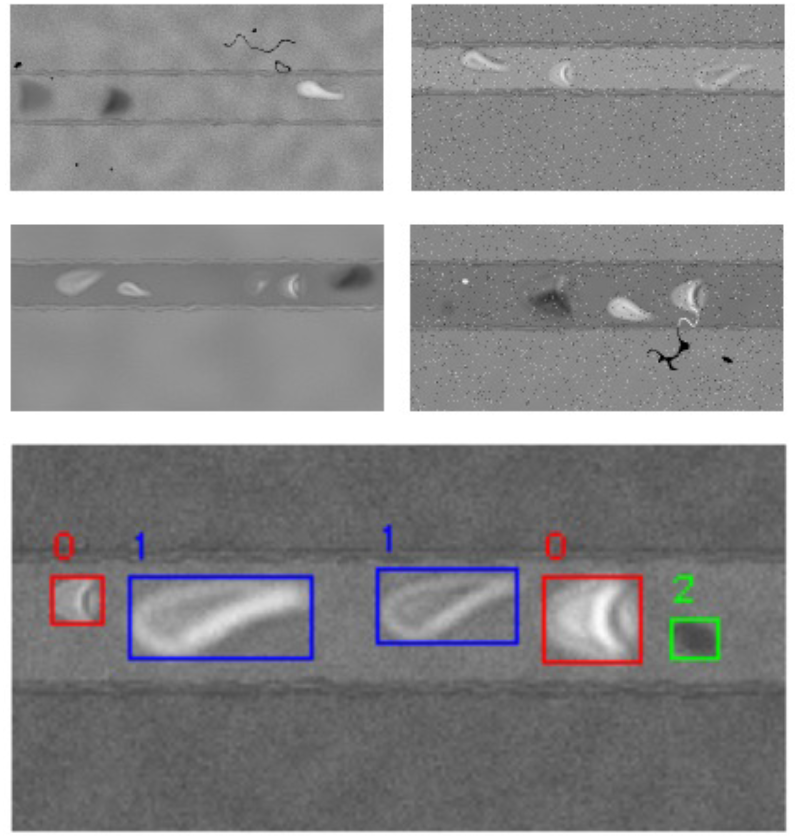
Example of generated images with the representation of microfluidic channels including multiple RBCs.

Both datasets — the one with **randomly placed cells, with or without cell superposition** and the one that **mimics microfluidic channels** — are generated in two distinct versions to evaluate the model’s generalization capabilities:

- **Two-Class Version:** This version classifies cells into two primary classes: parachute and slipper,
- **Three-Class Version:** Expanding on the two-class version, this iteration includes an additional class, classifying cells as parachute, slipper, or other.

### C. YOLO performance

YOLO is trained on the dataset consisting of images extracted from [15], [23], and [18], and tested on images extracted from videos [23] and [33]. The YOLOv11s model is trained for 20 epochs using the ‘detect’ task, with a batch size of 16 and an image size of 200 *×* 200 pixels. When testing on [23], the test images include cells along with their surrounding background, as shown in Figure 10a. Since the original dataset does not include labels (parachute or slipper), we implement a label inference algorithm similar to that described in Section III B 2. However, some cells are misclassified because the filters are too restrictive: croissants and slippers are labeled correctly, but cells classified as *other* may in fact be croissants or slippers. Similarly, when testing on [33], the test images include a cell and its surrounding background, as in the original dataset (Figure 10b). In this case, our labeling algorithm does not provide acceptable classifications; therefore, all cells are labeled as *other*. Thus, the objectives of this section depend on the specific test dataset:

**FIG. 10:**
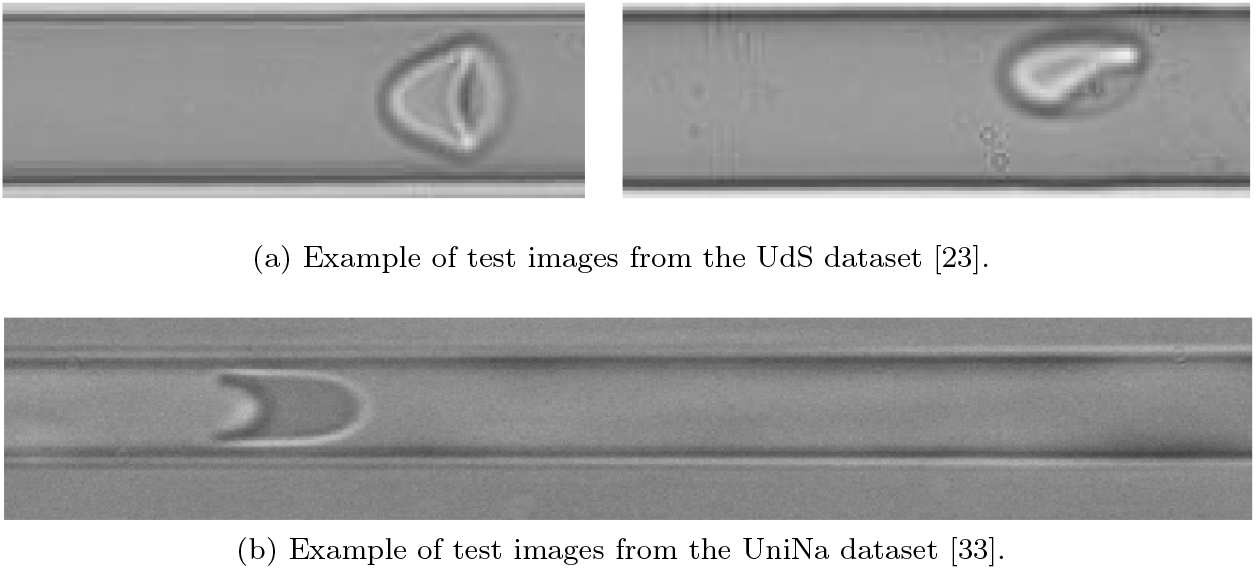
Examples of images on which the models were tested.

- For the UdS dataset [23], the focus should be on how accurately the model classifies croissants and slippers,
- For the UniNa dataset [33], since the images differ significantly from the training data (in terms of cell shapes and color profiles), the goal of testing on this dataset is to evaluate whether the model can effectively distinguish cells from the background.

#### 1. The UdS dataset

The first results analyzed are those obtained by testing the model on UdS videos. Considering the confusion matrix for the model trained on the dataset with randomly placed cells, allowing superposition, and classified into three classes (Figure 11a), the model demonstrates excellent performance in classifying the three cell shapes. Specifically, the parachute class exhibits an outstanding true positive rate (recall) of 0.99, indicating that nearly all actual parachute cells were correctly identified. While performance for the other two classes is generally good, a notable portion of actual slipper cells are misclassified as *other* (0.14), and conversely, some *other* cells are misclassified as slippers (0.34). This indicates a degree of morphological ambiguity or feature overlap between slipper and other cells. Furthermore, a substantial fraction (0.22) of *other* cells are misclassified as background, suggesting that some cells labeled as *other* are very small, faint, or poorly defined, making them difficult to distinguish from non-cellular regions. Analysis of background misclassifications also suggests that features learned from isolated cells may sometimes be erroneously triggered by noise or non-cellular elements in the video frames. Overall, these results highlight the model’s strong capability when trained on data that realistically represents the target environment. In particular, including superimposed cells appears crucial for achieving robust performance on complex video data.

**FIG. 11:**
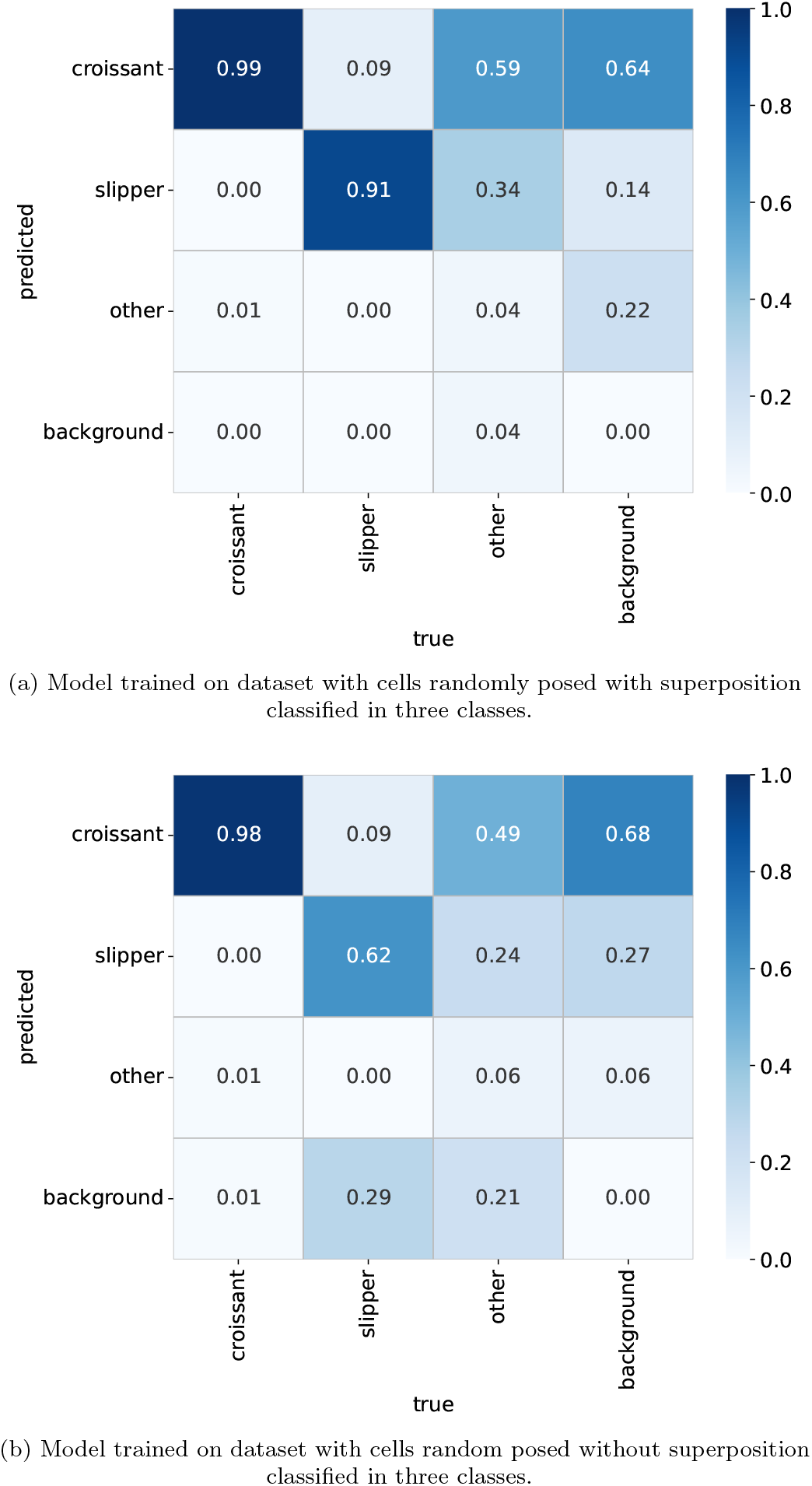
Confusion matrices evaluated on videos provided by the UdS.

Regarding the confusion matrix for the model trained on a dataset with randomly placed cells without superposition, classified into three classes (Figure 11b), the performance decreases significantly, particularly for the slipper and *other* categories. The parachute class maintains a high recall of 0.98, reaffirming its robust detectability regardless of whether superposition is present in the training data. However, the slipper class shows a notable drop in recall to 0.62, and *other* cells are frequently misclassified, predominantly as parachutes.

The comparison between the two confusion matrices highlights the significant impact of training data characteristics, specifically the presence or absence of cell superposition on model performance. Both models maintain exceptional performance for the parachute class, suggesting that parachute cells possess highly distinctive features that are robust to variations in superposition within the training data. Notably, the model trained with superposition demonstrates markedly better generalization, particularly for the slipper and *other* classes. The *other* class is inherently heterogeneous, encompassing various cell shapes that do not fit into parachute or slipper categories, which makes it more challenging for the model to learn a consistent set of features. Another contributing factor to misclassification may be that the ground truth annotations, generated algorithmically, are not entirely precise. In such cases, the model’s apparent “misclassifications” could reflect ambiguities or inaccuracies in the algorithmic ground truth rather than true failures in cell identification.

Due to the observed misclassification between the slipper and *other* classes, the same model was retrained to predict only two classes: parachute and slipper. The resulting confusion matrix is shown in Figure 12. Overall, performance is strong for both primary cell types. The parachute class again demonstrates superior detection, achieving a high recall of 0.95, with only a minor portion (0.07) of true parachute cells misclassified as slipper, indicating very high precision. The slipper class also performs reasonably well, with a recall of 0.55. However, compared to the three-class models, the misclassification rate between the two main cell types is higher, reflecting the inherent challenge in distinguishing slipper cells from parachutes when *other* cells are excluded.

**FIG. 12:**
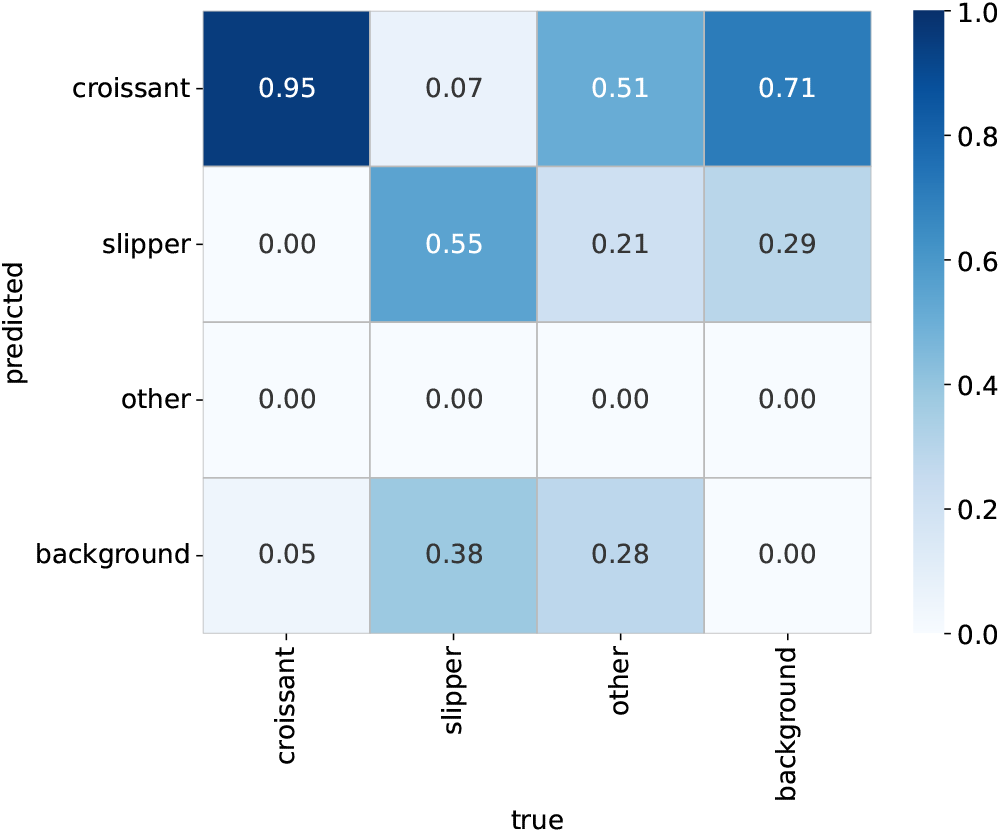
Confusion matrix evaluated on UdS videos using a model trained on a dataset of randomly posed cells with superposition, classified into two classes.

When contrasting the performance of the three-class models with the two-class model, several key observations emerge. First, the parachute class consistently achieves outstanding recall across all scenarios, regardless of the number of classes or the presence of cell superposition in the training data. This reinforces the hypothesis that its distinctive morphology makes it inherently easy to detect, serving as a reliable baseline metric for model capability. Second, simplifying the problem to two classes does not improve the performance of the slipper class. In fact, in the two-class model, a higher proportion of true slipper cells are misclassified as background (0.38) compared to the three-class models, both with and without superposition. This suggests that removing the *other* class does not inherently make slipper cells more distinguishable from the background. On the contrary, forcing the model to make a binary decision between ‘slipper’ and ‘background’ (or ‘parachute’ and ‘background’) may increase the likelihood that slipper cells are mistaken for non-cellular regions.

Finally, YOLO is trained on the dataset containing micro-channels, and the results are summarized in the confusion matrix shown in Figure 13. Both the croissant and slipper classes exhibit high true positive rates of 0.99 and 0.97, respectively, demonstrating strong detection capabilities. On the downside, the model is unable to distinguish any cells beyond the parachute and slipper categories, highlighting a limitation in handling additional or heterogeneous cell types. Comparing these results to the models trained on randomly posed cells, the consistent, near-perfect recall for the parachute class across all training scenarios (randomly posed with or without superposition, and microchannels) strongly reinforces the conclusion that its distinctive morphology makes it inherently easy for the model to learn and detect, serving as a robust baseline for model performance. For the slipper class, performance improves significantly in the micro-channel context compared to the randomly posed dataset without superposition (0.97 recall vs. 0.62), and is comparable to the model trained on superposed random cells classified into three classes. Training on microchannel data enables the model to excel at identifying the two most prevalent or clearly defined cell types in that specific environment by effectively collapsing the ‘other’ category into parachute, slipper, or background. In contrast, models trained on randomly posed cells, particularly with superposition, attempt a broader three-class classification. While they achieve decent performance for the slipper and other classes when superposition is included, they still experience notable misclassifications between these ambiguous categories and with the background. The consistent difficulties in classifying the ‘other’ class across both microchannel and randomly posed datasets highlight an inherent trade-off in model design: balancing broad generalization with the need for specialized, high-accuracy detection of well-defined classes within a specific context.

**FIG. 13:**
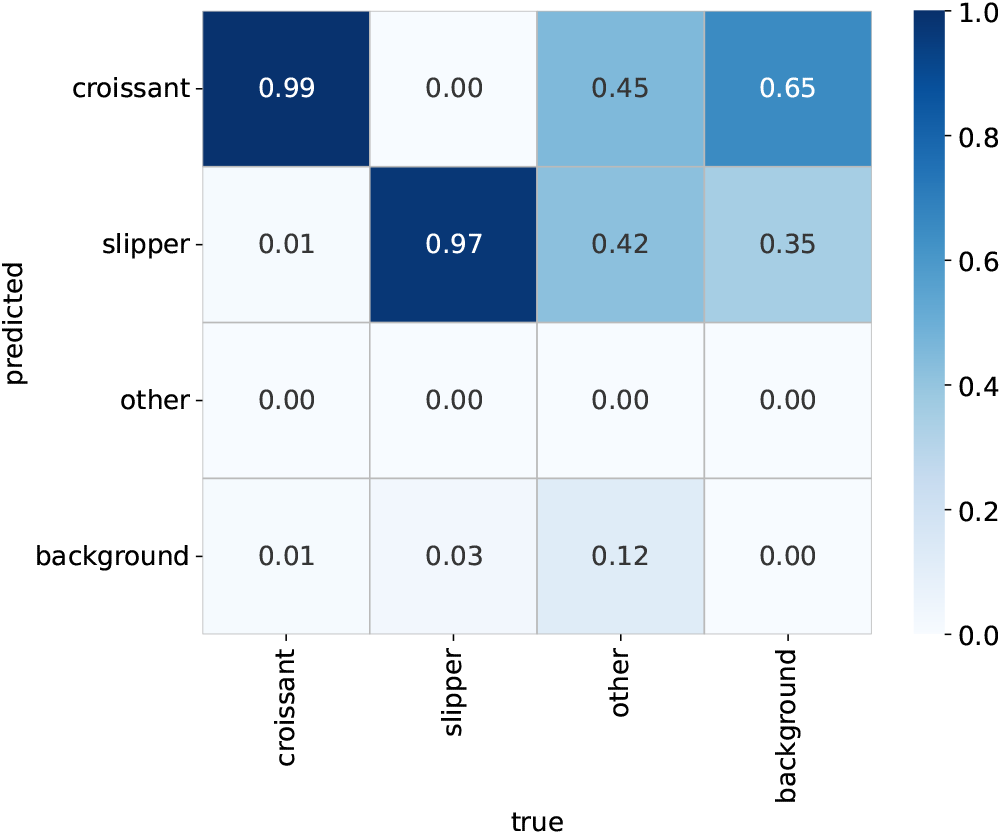
Normalized confusion matrix on UdS videos using a model trained on a dataset with microchannel structures, classified into three classes.

#### 2. The UniNa dataset

The YOLO framework is also tested on a separate dataset, whose images differ significantly from the training data in terms of cell shapes and color profiles. As shown in Figure 10b, this dataset contains elongated cells with cores that blend into the background. Therefore, the primary task in this evaluation is detecting cells relative to the background. Ground truth labels are unavailable for this dataset, and they cannot be inferred with sufficient precision. Consequently, all cells in these test experiments are labeled as ‘*other* ‘.

The first results come from the model trained on randomly placed cells that allows overlaps. The corresponding confusion matrix is shown in Figure 14a. It should be noted that the ground truth columns for *croissants* (parachutes) and *slippers* are empty. This is a direct consequence of the labeling policy for this dataset, where all cells are labeled as *other*. While the model predictions are available, the ground truth categories are not, and this holds for all test experiments on this dataset. From confusion matrix 14a, it can be observed that the model detected 79% of all cells present in the images (0.25 + 0.52 + 0.02), missing the remaining 21%. A substantial portion of the predicted cells corresponded to background regions: out of the 428 predicted cells (sum of croissant, slipper, and other rows in the non-normalized confusion matrix), exactly 50% were actually background portions of the images. This indicates that the model interprets background features as cells, i.e., it exhibits a high false positive rate. These results reflect the significant discrepancy between the test images and the training images, highlighting that the model was unable to generalize effectively to this unseen dataset.

**FIG. 14:**
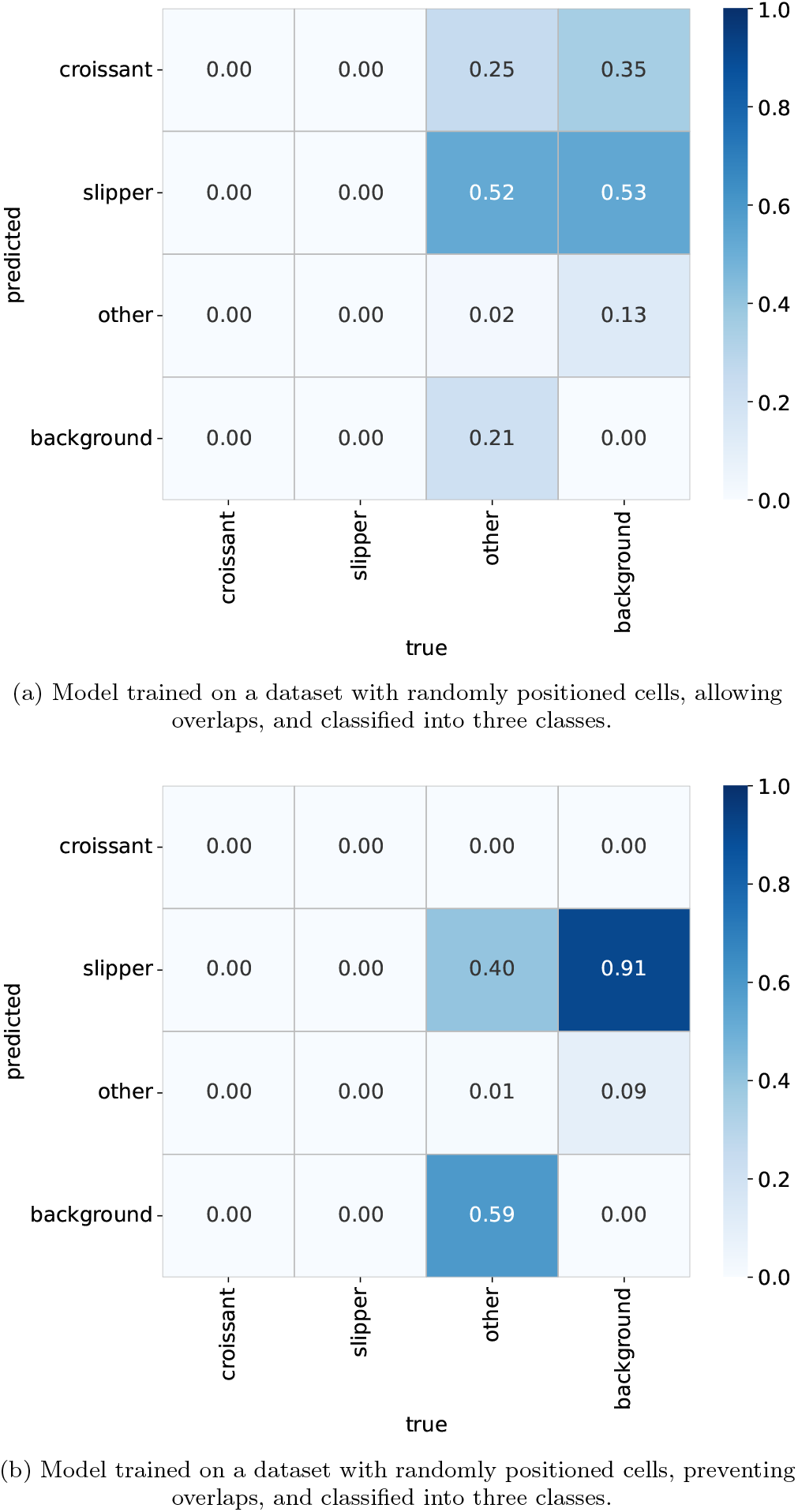
Normalized confusion matrix on UniNa videos for models trained with randomly placed cells and three-class classification. The distinguishing factor is the presence of overlaps.

To test whether the lack of generalization stems from the overlap of cells in the training data, we evaluated the model trained with randomly positioned cells without overlap. The resulting confusion matrix is shown in Figure 14b. An unexpected finding is that this model does not predict any parachutes, even though the test images contain them.

Additionally, it misses more cells overall, with 160 undetected cells—a 180% increase compared to the overlapping-cell model. However, it misinterprets the background less: only 33 background regions were incorrectly detected as cells, compared to 214 in the previous model. In conclusion, this model “hallucinates” less, meaning it interprets background features as cells less frequently, but at the cost of missing more real cells. This suggests it is more robust to novel backgrounds while being less sensitive to previously unseen cell objects.

When trained on only two classes, the model performs substantially worse: 83% of cells are classified as background, and 100% of background regions are misclassified as slippers. This indicates that the model fails to generalize and accurately distinguish between cells and background, resulting in a high rate of missed cells and a complete misinterpretation of background regions as a specific cell type. These results suggest either significant overfitting or an inherent inability of the model to learn the relevant discriminative features when restricted to only two classes.

A similar trend is observed when the model is trained on the dataset mimicking microfluidic channels: 88% of cells are classified as background. This indicates that this model also exhibits a very high false negative rate, failing to correctly identify the majority of actual cells and misclassifying them as background, analogous to the previous case with randomly positioned cells.

Across all test experiments on this challenging dataset, a consistent trend emerges: despite its general capabilities, the YOLO framework struggles significantly to generalize when there is a marked discrepancy between training and test image characteristics. Models trained on randomly placed cells exhibited differing trade-offs depending on the presence of overlap. The overlap model (Figure 14a) detected more cells (79%) but suffered a high false positive rate, misinterpreting 50% of predictions as background. In contrast, the non-overlap model (Figure 14b) reduced background misinterpretation but missed a substantially higher proportion of actual cells. When the number of classes was reduced to two, or when the training data mimicked microfluidic channels (Figures 15 and 16), performance deteriorated dramatically, with 83% and 88% of actual cells, respectively, misclassified as background. Collectively, these results underscore the models’ difficulty in adapting to novel cell shapes and blending colors in the test dataset, often resulting in high false-negative rates and, in some cases, severe false-positive rates where background features were misinterpreted as cells. This highlights the critical importance of training data fidelity for generalization to unseen, disparate image characteristics.

**FIG. 15:**
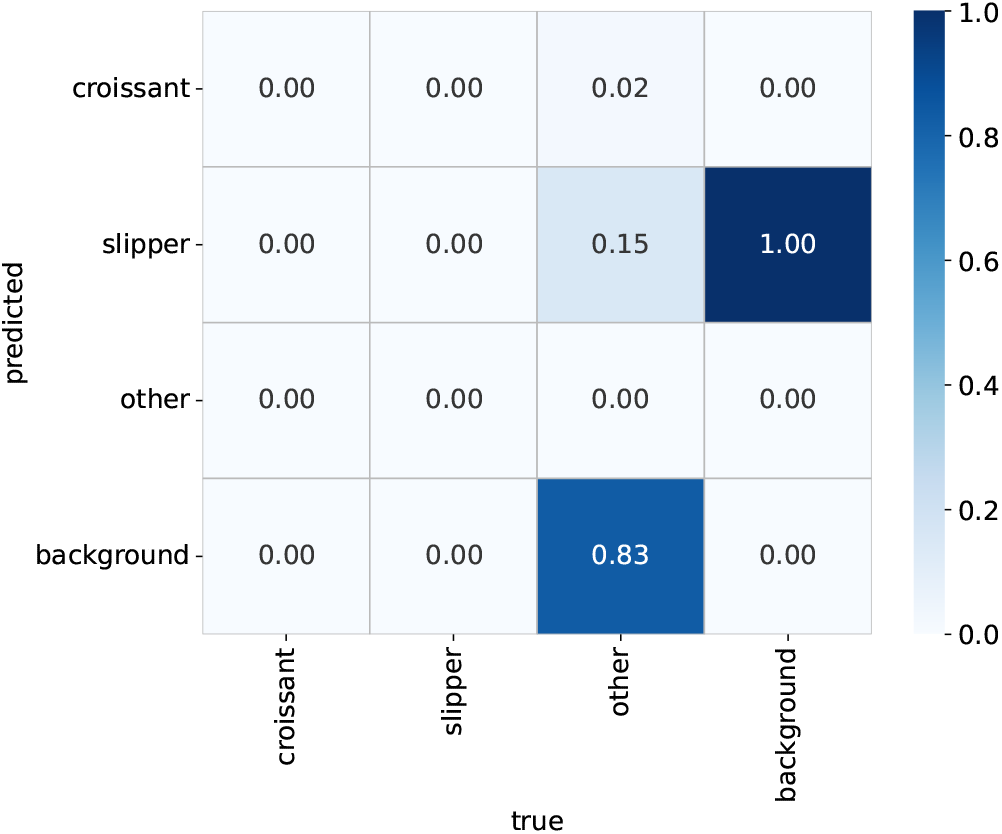
Normalized confusion matrix on the UniNa dataset for a model trained on randomly posed cells with superposition, performing two-class classification.

**FIG. 16:**
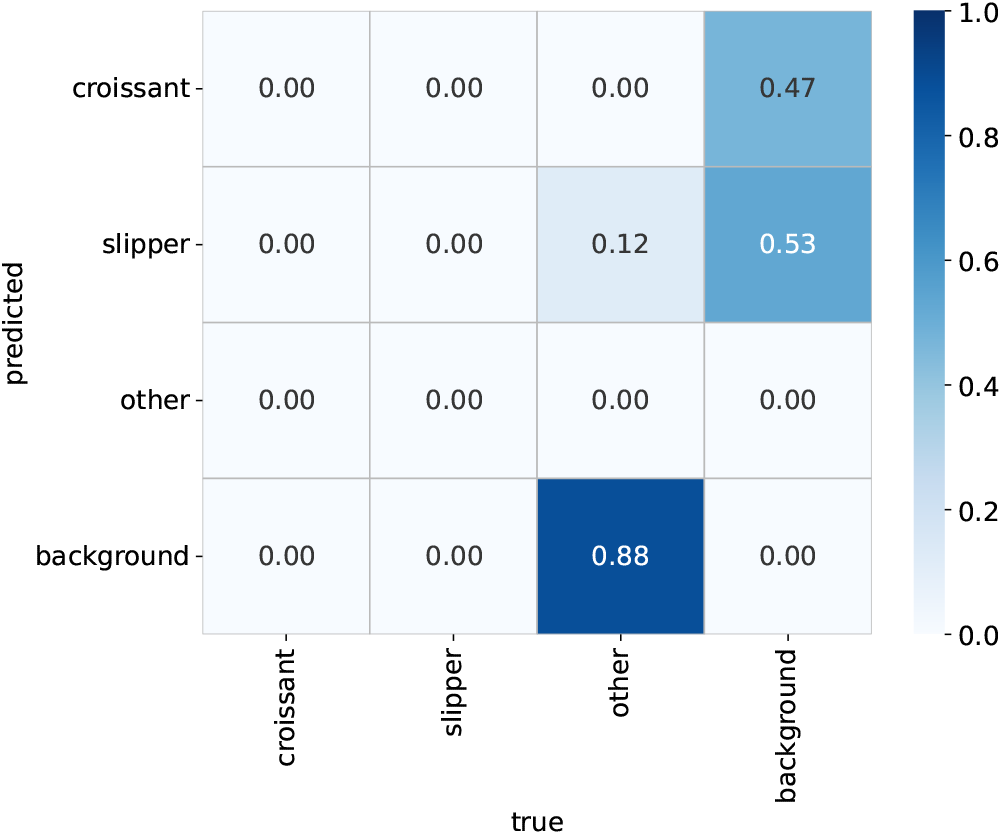
Normalized confusion matrix on the UniNa dataset: model trained on a dataset with microchannel structures (no overlaps), classified into three classes.

The comprehensive evaluation of the YOLO framework across two distinct datasets, UdS and UniNa, provides key insights into its performance, particularly regarding generalization and the influence of training data characteristics. On the UdS dataset, focused on accurate classification of cell types, the ‘parachute’ class consistently exhibited exceptional detectability across all training scenarios, including varying degrees of cell superposition and task simplification. This underscores the distinct morphological features of parachute cells, which make them robustly recognizable by the model. In contrast, performance for slipper and other classes was more sensitive to the training data: including cell superposition significantly improved generalization to these more ambiguous categories. Notably, simplifying the classification task to two classes did not universally enhance performance; it sometimes increased the misclassification of slipper cells as background, indicating that reducing class complexity alone does not guarantee better feature learning. The model trained on microchannel-mimicking data achieved high accuracy for croissant and slipper cells within that specific context, highlighting the advantage of specialized training for well-defined classes in targeted environments.

In stark contrast, tests on the UniNa dataset—characterized by markedly different cell shapes and color profiles blending with the background—consistently revealed the YOLO model’s struggle with generalization. Across all training data configurations, whether with or without overlap, two-class setups, or microchannel-mimicking datasets, the models exhibited alarmingly high false negative rates, failing to detect a substantial majority of actual cells.

Additionally, varying degrees of false positives were observed, with background features frequently misinterpreted as cells, highlighting the model’s limited ability to distinguish true cells from challenging backgrounds.

These results collectively underscore that, while YOLO can achieve high performance on datasets similar to its training data or for classes with highly distinctive features, its generalization ability is severely compromised when confronted with significant domain shifts, particularly in cell morphology and background characteristics. The study thus emphasizes the critical importance of diverse and representative training data that closely mirrors real-world conditions, illustrating the fundamental trade-off between achieving specialized, high-accuracy detection and maintaining robust generalization across highly disparate visual domains.

## IV. CONCLUSIONS

This study investigated the detection and classification of red blood cell morphologies, specifically “slipper” and “parachute” shapes, using YOLOv11, with a focus on addressing data sparsity and bias through tailored dataset construction. Single-cell datasets were created using segmentation models (U-Net, Cellpose) and image processing techniques to produce standardized, well-centered RBC images. These images, while limited for direct object detection, provided precise labeling and normalization, forming a solid basis for data augmentation.

To improve generalization, multiple-cell datasets were generated via cut-and-paste augmentation, producing three variants: allowing overlap, prohibiting overlap, and mimicking microfluidic channels. These datasets diversified training material, exposing the model to variations in cell arrangement, density, and background characteristics. Correlation analysis confirmed independence of generated image characteristics, ensuring robust diversity. On the UdS dataset, YOLOv11 achieved consistently high recall for the distinct “parachute” class. Inclusion of overlapping cells improved classification of the more ambiguous “slipper” and “other” categories. Simplifying to two classes did not always enhance slipper detection and sometimes increased false negatives. In contrast, on the UniNa dataset, characterized by different cell shapes and background blending, the model struggled with generalization. High false negative rates were observed, with many real cells missed, and false positives occurred due to background misinterpretation.

Overall, YOLOv11 performs well on data similar to its training set but is limited when faced with substantial domain shifts. This highlights the need for diverse, representative training data to achieve robust performance in real-world microscopy applications. The code and data sets are available on GitHub [1].

## V. PERSPECTIVES

This study introduced a novel dataset construction method for RBC detection and classification, but several avenues remain to enhance robustness, generalization, and clinical applicability. Advanced data augmentation techniques could further improve model performance. Future work might explore synthetic data generation approaches, including image in-painting to remove and refill cells while preserving realistic microfluidic backgrounds. This would enable controlled cell densities and rare morphological states difficult to achieve experimentally. Parametric manipulation of cell characteristics, such as illumination, contrast, and optical distortions, could also improve robustness across varying imaging conditions. Generative models, like GANs or diffusion models, could create large-scale photorealistic RBC datasets with controlled morphology, reducing the need for labor-intensive manual segmentation. Improving generalizability requires more diverse datasets from multiple institutions, covering varied microfluidic designs, imaging systems, preparation protocols, and population demographics. Including additional cell types with proper labeling would further enhance model accuracy in realistic biological samples. Finally, while YOLOv11 performed well in current tasks, future studies could explore alternative object detection frameworks, such as Vision Transformers, to further advance RBC detection and classification.

## Acknowledgment

The authors would like to thank Prof. Giovanna Tomaiuolo and Prof. Stefano Guido of the Università degli Studi di Napoli Federico II; Prof. Christian Wagner, Prof. Lars Kästner, and Dr. Mohammed Nouaman of the Universität des Saarlandes; and Prof. Ye Ai of the Singapore University of Technology and Design for kindly sharing their datasets and granting permission for their use. The authors also wish to thank Dr. Imad Rida, Ms. Alaa Bou Orm, and Prof. Amine Nait-Ali for their valuable support and for providing the opportunity to present this work at BioSMART—the 6th International Conference on Bioengineering for Smart Technologies.

## References

[1] A. Amrani, I. Caridi, and B. Kaoui, Detection and Classification of Red Blood Cells under Flow, GitHub repository (2025). https://github.com/4l3x4ndre/Detection-and-Classification-of-Red-Blood-Cells-under-flow

[2] S. Ben-David, J. Blitzer, K. Crammer, A. Kulesza, F. Pereira, and J. Wortman Vaughan, A theory of learning from different domains, Machine Learning, vol. 79, pp. 151–175, 2010, 10.1007/s10994-009-5152-4

[3] BioStudies, BioStudies - EMBL-EBI, https://www.ebi.ac.uk/biostudies/, accessed May 6, 2025.

[4] J. Brown, A. Nguyen, and N. Raj, Effect of Camera Choice on Image-Classification Inference, Applied Sciences, vol. 15, no. 1, p. 246, 2024, 10.3390/app15010246

[5] A. K. Dasanna, J. Mauer, G. Gompper, and D. A. Fedosov, Importance of viscosity contrast for the motion of erythrocytes in microcapillaries, Frontiers in Physics, Sec. Biophysics, vol. 9, 666913, 2021, 10.3389/fphy.2021.666913

[6] D. Dwibedi, I. Misra, and M. Hebert, Cut, Paste and Learn: Surprisingly Easy Synthesis for Instance Detection, arXiv:1708.01642, 2017, 10.48550/arxiv.1708.01642

[7] M. Ester, H.-P. Kriegel, J. Sander, and X. Xu, A density-based algorithm for discovering clusters in large spatial databases with noise, Proceedings of the Second International Conference on Knowledge Discovery and Data Mining (KDD-96). AAAI Press. pp. 226–231, 1996, ISBN 1-57735-004-9, https://cdn.aaai.org/KDD/1996/KDD96-037.pdf

[8] Figshare, figshare - Where research data lives, https://figshare.com/, accessed May 6, 2025.

[9] Y. Ganin and V. Lempitsky, Unsupervised Domain Adaptation by Backpropagation, arXiv:1409.7495, 2015, 10.48550/arxiv.1409.7495

[10] GitHub, GitHub: Where the world builds software · GitHub, https://github.com/, accessed May 6, 2025.

[11] N. Jegham, C. Y. Koh, M. Abdelatti, and A. Hendawi, YOLO evolution: A comprehensive benchmark and architectural review of YOLOv12, YOLO11, and their previous versions, arXiv:2411.00201v4, 2025, 10.48550/arXiv.2411.00201

[12] B. Kaoui, G. Biros, and C. Misbah, Why do red blood cells have asymmetric shapes even in a symmetric flow?, Physical Review Letters, vol. 103, 188101, 2009, 10.1103/PhysRevLett.103.188101

[13] B. Kaoui, N. Tahiri, T. Biben, H. Ez-Zahraouy, A. Benyoussef, G. Biros, and C. Misbah, Complexity of vesicle microcirculation, Physical Review E 84, 041906, 2011, 10.1103/PhysRevE.84.041906

[14] R. Khanam and M. Hussain, YOLOv11: An overview of the key architectural enhancements, arXiv:2410.17725, 2024, 10.48550/arXiv.2410.17725

[15] A. Kihm, L. Kaestner, C. Wagner, and S. Quint, Classification of red blood cell shapes in flow using outlier tolerant machine learning, PLoS Computational Biology, vol. 14, no. 6, e1006278, 2018, 10.1371/journal.pcbi.1006278

[16] Kaggle, Kaggle: Your Machine Learning and Data Science Community, https://www.kaggle.com/, accessed May 6, 2025.

[17] K. T. Navya, K. Prasad, and B. M. K. Singh, Analysis of red blood cells from peripheral blood smear images for anemia detection: a methodological review, Medical and Biological Engineering and Computing, Volume 60, pages 2445–2462, 2022, 10.1007/s11517-022-02614-z

[18] M. Liang, J. Zhong, C. S. Shannon, R. Agrawal, and Y. Ai, Intelligent image-based deformability assessment of red blood cells via dynamic shape classification, Sensors and Actuators B: Chemical, vol. 401, 135056, 2024, 10.1016/j.snb.2023.135056

[19] Z. Liu, T. Lian, J. Farrell, and B. A. Wandell, Neural Network Generalization: The Impact of Camera Parameters, IEEE Access, vol. 8, pp. 10443–10454, 2020, 10.1109/access.2020.2965089

[20] milesial, Pytorch-UNet: UNet implementation in PyTorch, GitHub repository, 2018–2025. Available at: https://github.com/milesial/Pytorch-UNet (accessed 2025-05-08).

[21] NIH Data Commons, NIH Data Commons, https://commonfund.nih.gov/datacommons, accessed May 6, 2025.

[22] H. Noguchi and G. Gompper, Shape transitions of fluid vesicles and red blood cells in capillary flows, Proceedings of the National Academy of Sciences of the United States of America, vol. 102, no. 40, pp. 14159–14164, 2005, 10.1073/pnas.0504243102

[23] M. Nouaman, A. Darras, T. John, G. Simionato, M. A. E. Rab, R. van Wijk, M. W. Laschke, L. Kaestner, C. Wagner, and S. M. Recktenwald, Effect of cell age and membrane rigidity on red blood cell shape in capillary flow, Cells, vol. 12, no. 11, 1529, 2023, 10.3390/cells12111529

[24] F. Pedregosa, G. Varoquaux, A. Gramfort, V. Michel, B. Thirion, O. Grisel, M. Blondel, A. Prettenhofer, R. Weiss, V. Dubourg, J. Vanderplas, M. Joly, B. Holt, G. Andrianov, M. Machineni, M. Brett, A. Mensch, J. Vincent, A. Le Borgne, E. Ghayes, C. Joly, B. Moreau, F. Gauthier, G. Louppe, M. Bromberg, A. Buchacher, J. Pichon, A. Ballan, A. Salmon, A. Fromont, K. Bernard, Y. Vendier, V. Gramfort, L. Buitinck, A. Lacoste, G. Jewson, A. Estimé, P. Massich, J. Perrotin, F. Mueller, Z. Harchaoui, M. I. Jordan, L. Buitinck, G. Louppe, A. Prettenhofer, and R. Weiss, Scikit-learn: Machine learning in Python, Journal of Machine Learning Research, vol. 12, pp. 2825–2830, 2011, 10.48550/arXiv.1201.0490

[25] D. T. Rademaker et al., Quantifying the deformability of malaria-infected red blood cells using deep learning trained on synthetic cells, iScience, vol. 26, no. 12, 108542, 2023, 10.1016/j.isci.2023.108542

[26] S. M. Recktenwald, K. Graessel, F. M. Maurer, T. John, S. Gekle, and C. Wagner, Red blood cell shape transitions and dynamics in time-dependent capillary flows, Biophysical Journal, vol. 121, no. 1, pp. 23–36, 2022, 10.1016/j.bpj.2021.12.009

[27] J. Redmon, S. K. Divvala, R. B. Girshick, and A. Farhadi, YOLO: Real-time object detection, arXiv:1506.02640, 2016, 10.48550/arXiv.1506.02640

[28] ResearchGate GmbH, ResearchGate: Discover and share research, https://www.researchgate.net/, accessed May 10, 2025.

[29] O. Ronneberger, P. Fischer, and T. Brox, U-Net: Convolutional Networks for Biomedical Image Segmentation, arXiv:1505.04597, 2015. 10.48550/arXiv.1505.04597

[30] C. Stringer, T. Wang, M. Michaelos, and M. Pachitariu, Cellpose: a generalist algorithm for cellular segmentation, Nature Methods, vol. 18, pp. 100–106, 2021, 10.1038/s41592-020-01018-x

[31] N. Takeishi, H. Ito, M. Kaneko, and S. Wada, Deformation of a red blood cell in a narrow rectangular microchannel, Micromachines, vol. 10, no. 3, p. 199, 2019, 10.3390/mi10030199

[32] N. Takeishi, H. Yamashita, T. Omori, N. Yokoyama, and M. Sugihara-Seki, Axial and nonaxial migration of red blood cells in a microtube, Micromachines, vol. 12, no. 10, 1162, 2021, 10.3390/mi12101162

[33] G. Tomaiuolo, M. Simeone, V. Martinelli, B. Rotoli and S. Guido, Red blood cell deformation in microconfined flow, Soft Matter, vol. 5, pp. 3736—3740, 2009, 10.1039/b904584h

[34] G. Tomaiuolo and S. Guido, Start-up shape dynamics of red blood cells in microcapillary flow, Microvascular Research, vol. 82, no. 1, pp. 35–41, 2011, 10.1016/j.mvr.2011.03.004

[35] A. Torralba and A. A. Efros, Unbiased look at dataset bias, CVPR 2011, Colorado Springs, CO, USA, pp. 1521–1528, 2011, 10.1109/CVPR.2011.5995347

[36] S. van der Walt, J. L. Schönberger, J. Nunez-Iglesias, F. Boulogne, J. D. Warner, N. Yager, E. Gouillart, T. Yu, and the scikit-image contributors, scikit-image: image processing in Python, PeerJ, vol. 2, e453, 2014, 10.7717/peerj.453

[37] W-H. Yun, T. Kim, J. Lee, J. Kim, and J. Kim, Cut-and-Paste Dataset Generation for Balancing Domain Gaps in Object Instance Detection, IEEE Access, vol. 9, pp. 14319–14329, 2021, 10.1109/access.2021.3051964

[38] Zenodo, Zenodo: Research. Shared., https://zenodo.org/, accessed May 6, 2025.

